# Unlocking Scalable Ligand Residence Time Predictions with Koffee Unbinding Kinetics Simulations

**DOI:** 10.1101/2025.11.06.686759

**Authors:** Niels Kristian Madsen, Robert M. Ziolek, Daniel Kongsgaard, Christian Flohr Nielsen, Anders Dyhr Nørløv, Daniela Dolciami, Joshua R. Sacher, Klaus Michelsen, Michael G. Acker, Nils Anton Berglund, Mikael H. Christensen, Allan Grønlund, Lise Husted, David E. Gloriam, Albert J. Kooistra, Nikolaj Thomas Zinner

**Author notes:** N.K.M. and R.M.Z. contributed equally.

## Abstract

A great number of drug discovery programs fail due to poor *in vivo* efficacy and ADMET liabilities. On- and off-target ligand residence times can act as important drivers of these problems. While modern experimental techniques have made measuring compound kinetics data more routine, there is a lack of accurate, high-throughput simulation techniques to guide compound prioritization by residence time. In this work, we introduce *Koffee*™ *Unbinding Kinetics* as a solution to the hitherto unanswered problem of scalable ligand-protein residence time prediction. By bypassing conventional approaches based on molecular dynamics simulations, *Koffee Unbinding Kinetics* performs physics-based residence time screening at the atomistic level in ≈ 1 GPU minute per complex using inexpensive hardware, a speed-up of at least 3 − 5 orders of magnitude compared to current state-of-the-art simulation approaches. *Koffee Unbinding Kinetics* can enhance compound selection to mitigate costly future program failures by adding fast, predictive residence time simulations to early-stage computational drug discovery pipelines.

**TOC Graphic**

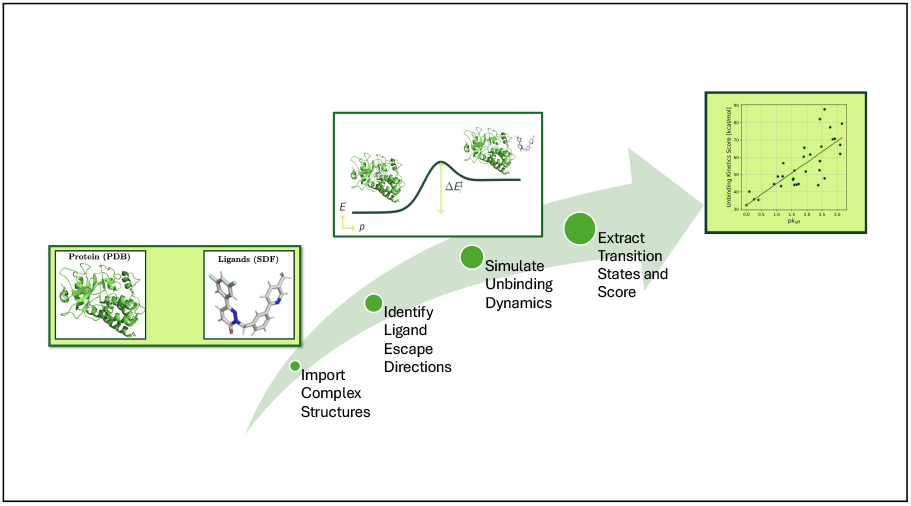

## Introduction

The kinetics of ligand-protein residence time (RT) has historically been overlooked by drug discovery scientists in favor of the equilibrium thermodynamics of binding affinity. ^1,2^ In more recent times, new experimental techniques have supported the growth of RT measurements in drug discovery, while modern computer simulation techniques have begun to complement these experimental capabilities. Careful consideration of ligand RTs may help mitigate efficacy and ADMET issues in late-stage drug discovery,^3^ which are both major causes of clinical-stage program failure. ^4^ RTs offer a different perspective to binding affinity for compound optimization. Intriguingly, the relationship between RT and ligand efficacy is still to be fully understood and in some instances, RT is a better predictor of functional efficacy than binding affinity.^5^ Several techniques exist to measure ligand binding kinetics experimentally, including radioligand competition binding using radiolabeled ligands to measure ligand kinetics, time-resolved fluorescence energy transfer (that monitors energy transfer between a protein and ligand both labeled with fluorophores) and surface plasmon resonance (SPR) - a label-free approach using refractive index measurements.^6^ Since the ligand unbinding process and corresponding transition state ensemble cannot be examined directly by experiments due their inherently short lifetimes,^7^ experiments alone do not provide a means to optimize ligand-protein RTs by rational design. Instead, an incomparable level of mechanistic detail of fast molecular processes, such as ligand unbinding, can be obtained from modern computer simulation techniques.

Present state-of-the-art simulation tools to estimate ligand unbinding are based on molecular dynamics (MD) simulations, the so-called “computational microscope”.^8^ In recent years several new simulation approaches have been reported, powered by advances in algorithms and improved computational resources. ^6,9^ The timescales of ligand unbinding typically range from (sub)-seconds to hours making direct simulation by MD infeasible given its inherent sampling limitations. To circumvent this fundamental limitation, several research groups have developed innovative enhanced sampling MD methods to simulate ligand unbinding with greater computational efficiency. Approaches include random accelerated MD (RAMD),^10^ scaled MD, ^11^ milestoning simulations,^12,13^ adiabatic-bias MD (ABMD),^14^ and metadynamics.^15^ A recent report has highlighted how two MD methods, RAMD and infrequent metadynamics, can be deployed in combination to provide quantitative ligand RTs.^16^ Furthermore, Serrano-Morrás and co-workers have recently explored the use of steered MD simulations (as part of the “dynamic undocking” procedure) to compute quasi-bound states, which highlights an interesting link between ligand RT and binding affinity. ^17^

While these MD-based methods indeed make ligand-protein RT prediction computationally tractable, they still have simulation timescales ranging from tens of nanoseconds to several microseconds per complex. The longer end of this scale corresponds to several days of GPU time per ligand-protein complex, making the majority of these methods incompatible with the rapid decision-making cycles inherent to modern drug discovery. Instead of augmenting MD simulations, themselves an inherently equilibrium methodology, we took a different approach when developing *Koffee*™ *Unbinding Kinetics*. We built an alternative simulation engine specifically to model the nonequilibrium ligand unbinding process. *Koffee Unbinding Kinetics* simulates the ligand-protein unbinding process in around a minute per complex on a single inexpensive GPU (NVIDIA T4). When considering the number of force evaluations required, each of our simulations is equivalent to approximately 10 − 100 ps of MD simulation. *Koffee Unbinding Kinetics* represents a computation speed increase (and computational cost savings) of at least 3 − 5 orders of magnitude compared to any existing MD-based alternatives. Our new approach delivers accurate, high-throughput RT predictions, making ligand RTs practically accessible to prospective computational drug discovery tasks.

## Methods

Existing simulation techniques to predict ligand-protein RTs are based on time-resolved MD simulations. Given the large energy barrier to ligand unbinding, spontaneous ligand-protein dissociation cannot be directly simulated. Therefore, enhanced sampling methods apply an external bias to the system before the final result is obtained by a reweighting scheme. In this work, we reformulate the ligand unbinding problem from a time-resolved simulation into an energy-resolved optimization process that aims to find low-energy unbinding pathways to model the ligand unbinding (dissociation) process and find a representative transition state as the maximum energy state along the unbinding pathway. The Arrhenius equation establishes the relationship between the ligand-protein residence time (RT) and the activation energy (Δ*E*^‡^) of the unbinding process,

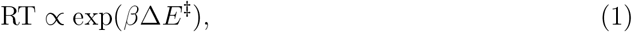

where the activation energy is simply defined as the difference between the bound state energy (*E*^bound^) and the transition state energy (*E*^‡^),

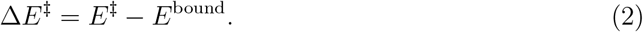

The challenging requirement for the simulations is therefore finding an unbinding pathway and transition state that faithfully models the real ligand-protein unbinding dynamics. We have developed a physical simulation algorithm for quickly simulating a number of nonequilibrium candidate unbinding pathways, 𝒫. The unbinding kinetics score is simply defined as the minimum activation energy of all computed unbinding pathways,

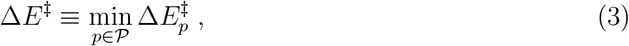

which is associated to the path *p* ∈ 𝒫. A schematic representation of ligand-protein unbinding is presented in Figure 1(a).

**Figure 1:**
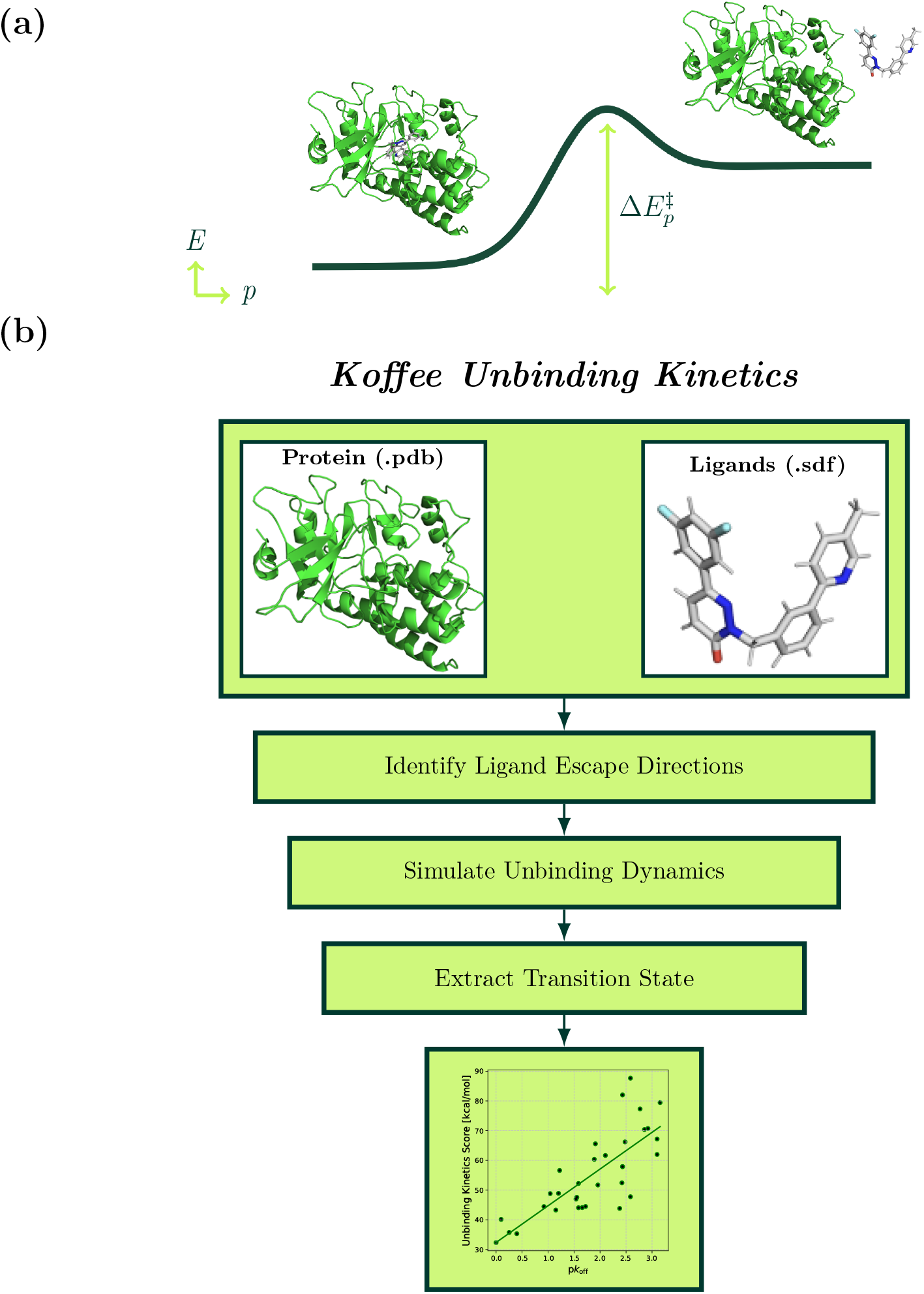
Schematic outline of the theory and methodology behind *Koffee Unbinding Kinetics*. Visualizations showing (a) a ligand-protein complex with a simplified representation of the energy (*E*) landscape (dark green curve) along the ligand unbinding path (*p*). The ligand-protein complex is initially bound in a low energy state (left hand side). Ligand unbinding dynamics move the system towards a transition state region (indicated by the peak in the energy curve) with an associated unbinding energy barrier 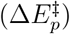. Full ligand-protein dissociation then occurs at the end of the path (right hand side). Molecules are not depicted to scale. (b) A schematic overview of the computational unbinding kinetics simulation method.

### Simulation Details

We use the Amber ff14SB force field^18^ to model proteins while ligands are modeled using the GAFF 2.11 force field.^19,20^ We compute ligand partial charges using the AM1-BCC semi-empirical method with the ligand input structures. Water is modeled using either the GBn or GBn2 implicit solvent models.^21,22^ We use the GBn model, which has wider chemical space coverage, in all cases except for the M_3_ mutants dataset (described later), where we use GBn2 as it was further tuned for use with peptides and proteins, in which the chemical differences lie in this dataset. In the case of the metalloprotein, Thermolysin, we employ the parameterization scheme originally reported by Tomić and co-workers to model zinc and its local coordination environment.^23^ When required, phosphorylated residues are described by the Amber phosaa14SB force field.^24^ Explicit water, whether present as crystal waters or bulk solvent, is not considered in the presented version of the algorithm and are automatically stripped from the input files. Furthermore, we do not consider generic binding cofactors or lipid membrane environments in the simulations presented in this work.

The unbinding kinetics simulation algorithm has been implemented using the OpenMM simulation engine (version 8.3.1). ^25^ A schematic representation of the computational unbinding kinetics method is depicted in Figure 1(b). The algorithm reads protein and ligand structures using the canonical .pdb and .sdf file types. System preparation and parameterization is then entirely automated, with no manual parameter tuning required. All simulation parameters, the implicit solvent model aside, are consistent in all of the results presented in this work, highlighting the robustness of our methodology. The simulation procedure begins with an initial energy minimization (maximum force tolerance 1.2 × 10^−2^ kcal mol^−1^Å^−1^) to embed the system into the force field energy landscape (no further structural refinement is performed). Subsequently, a number of candidate ligand escape directions are identified and the ligand unbinding simulation is initiated. Once full ligand dissociation is automatically detected, the corresponding transition state is extracted from the simulation trajectory for use in the final score. The score outputted is the raw system energy difference between the bound and transition states, with no rescaling or other mathematical manipulation. We developed the unbinding kinetics algorithm with a focus on making full use of GPU acceleration. All of the results in this work were computed using easily accessible NVIDIA T4 GPUs (one GPU per unbinding kinetics simulation), which are significantly less powerful than state-of-the-art hardware. Therefore, given that the majority of computational cost is associated with GPU time, it is reasonable to expect that GPU bottlenecking (and with it run time) could be significantly reduced by using more powerful, state-of-the-art GPU hardware.

### Datasets

Publicly available ligand unbinding kinetics data is far more scarce than corresponding datasets for binding affinity although efforts have been made to improve this situation. ^26,27^ We collected unbinding kinetics data from various sources to test our new methodology. The initial validation set consists of 7 different systems that contain between 10 and 86 ligands each. A summary of the datasets used in this work is provided in Table 1. Two datasets contain highly congeneric ligand series: eukaryotic translation initiation factor 4E (eIF4E) ^27,28^ and Muscarinic acetylcholine receptor M3 (M_3_).^29^ However, these are still challenging to simulate due to the large size of the surface binding ligands in eIF4E (which include a number of macrocycles) and the structural complexity of the M_3_ protein, a G protein-coupled receptor (GPCR). Two further datasets contain non-congeneric ligand series with varying formal charges for two kinases, focal adhesion kinase (FAK)^11,27^ and threonine tyrosine kinase (TTK).^30^ The FAK dataset is made up of 14 FAK ligands recently studied elsewhere by MD-based residence time simulations,^11^ which we supplemented with 19 ligand structures and corresponding *k*_off_ values obtained from PDBbind-koff-2020, ^27^ an unbinding kinetics-focused dataset published as part of the PDBbind database.^31^ Interestingly, the RTs of the former subset have a strong dependence on ligand molecular weight (*ρ*_MW_ = 0.79) while the FAK ligand set used in this work is only weakly correlated with respect to ligand molecular weight (*ρ*_MW_ = 0.22). HSP90 represents the largest single unbinding kinetics dataset in this work (*N* = 86) consisting of three main ligand series. ^10,27^ Thermolysin is included as we were interested to assess our approach on a metalloprotein.^27^ Finally, we included a set of M_3_ point mutants in complex with tiotropium (with single point mutations in the active site only) studied computationally elsewhere in the literature ^32^ to test the method’s applicability to this distinct task.

**Table 1:** Details of the datasets used in our internal validation of the unbinding kinetics method: number of ligands (**N**), average Tanimoto similarity of each ligand set (**T**) computed using iSIM,^35^ average percentage of atoms in the maximum common substructure (**MCS**%), the set of formal charges of ligands in each dataset ({**q**}), and references to original data sources.

| Target | Protein Family | N | T | MCS% | {q} | References |
| --- | --- | --- | --- | --- | --- | --- |
| eIF4E | Cap-binding protein | 10 | 0.90 | 72 | 2, 3, 4, 7 | 27,28 |
| FAK | Kinase | 33 | 0.53 | 35 | 0 | 11,27 |
| HSP90 | Molecular chaperone | 86 | 0.47 | 24 | -1, 0, 1 | 10,27 |
| M<sub>3</sub> | GPCR | 10 | 0.69 | 71 | 1 | 29 |
| M<sub>3</sub> mutants | GPCR | 13 | 1.00 | 100 | 1 | 32 |
| Thermolysin | Metalloenzyme | 17 | 0.90 | 87 | -2, -1 | 27 |
| TTK | Kinase | 10 | 0.45 | 28 | 0, 1 | 30 |

Except for when stated otherwise, complex structures extracted from PDBbind-koff-2020^27^ and Sinko *et al*.^11^ were used in their original form without modification. For TTK, we used PDBFixer^33^ to model missing loops and atoms and Protoss^34^ to asign protein and ligand protonation states, using default settings in both instances. We generated the M_3_ mutant structures by manually mutating the required residues with no further optimization, starting from the triotropium-M_3_ complex reported by Galvani *et al*. ^29^ For the Thermolysin dataset, we realigned all ligands using 5LWD as a template. Specific details of the exact composition of the amalgamated FAK and TTK datasets used in this work are provided in the Supporting Information, including PDB accession codes in the case of TTK. The datasets used in the collaboration between Kvantify and Delphia Therapeutics are proprietary to Delphia Therapeutics and are therefore not publicly available.

### Analysis Techniques

Ligand similarity was assessed as the average Tanimoto similarity between all ligands in each set (RDKit fingerprints) using the fast, linear-scaling algorithm, iSIM.^35^ Molecular visualizations and graphs were produced using the RDKit and Matplotlib, respectively.

## Results

### Applicability to Chemically Diverse Ligands and Binding Sites

Figure 2(a) shows results from our unbinding simulations for 6 different targets with chemically diverse ligands and binding sites. These datasets provide a broad initial test of the applicability of our approach. There are several potential complications in the eIF4E dataset: the surface binding, peptide-based ligands span a molecular weight range of 1361 to 2106 (far exceeding the 500 Da small molecule regime as described by Lipinski’s rule of five) with ligand net formal charges ranging from +2 to +7. We observe strong predictive performance for the eIF4E dataset despite these challenges and without including an explicit water treatment in our simulations. We also observe strong predictive ranking performance for the FAK dataset and note that the experimental off rates in this set have a particularly low correlation to ligand molecular weight as discussed in the Datasets section within Methods (see Table S1). We maintain good performance for M_3_ (a GPCR with its orthosteric site buried deep within the transmembrane domain), Thermolysin (a zinc-containing metalloprotein) and TTK (a highly chemically diverse ligand set, see Figure S2 for ligand chemical structures).

**Figure 2:**
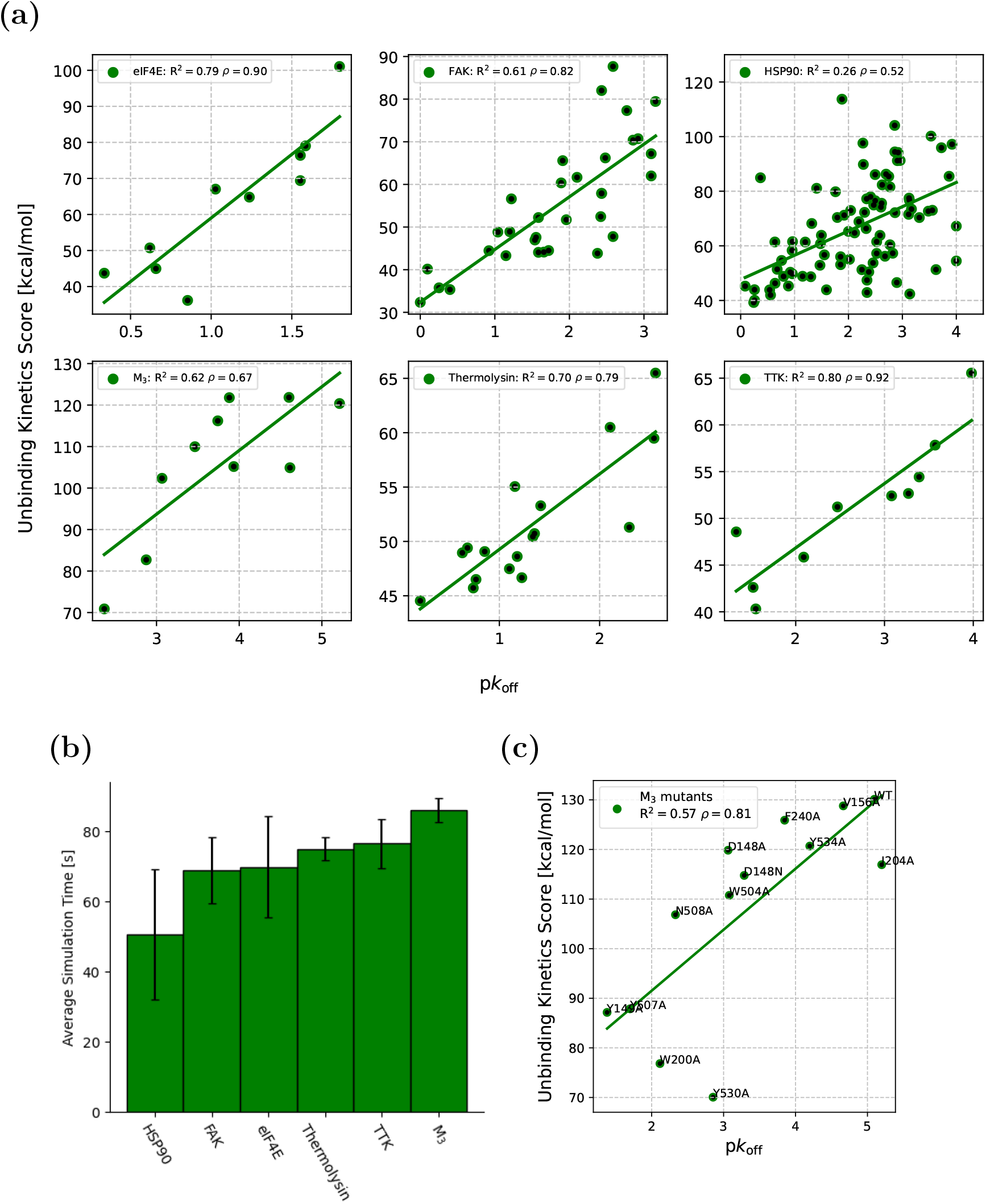
*Koffee Unbinding Kinetics* demonstrates broad applicability over chemically diverse ligands and binding sites. (a) Initial validation of the unbinding kinetics method on 6 different datasets. (b) Average simulation time per complex for the results reported in (a) where uncertainty bars represent 1 standard deviation. Simulations performed on a single NVIDIA T4 GPU. (c) Results of applying the unbinding kinetics method on the M_3_ mutants dataset.

The HSP90 dataset, a *n* = 70 subset of which was first modeled by Kokh and coworkers using *τ*-random acceleration MD (*τ*-RAMD) simulations, ^10^ is the largest publicly available set of structures for a single target with associated unbinding kinetics data (*n* = 86). The target has been used elsewhere as a testbed for developing and testing new unbinding kinetics simulation algorithms.^10,11,13^ Our performance on HSP90 in Figure 2(a) is poorer than the baseline ranking performance obtained from ligand molecular weight (*ρ*_MW_ = 0.76). This particularly strong influence of ligand molecular weight upon HSP90 ligand kinetics has been discussed elsewhere previously.^36^ While our ranking performance is reasonable, the linear correlation is more strongly impacted by a small number of outliers. We note for reference that when considering the entire set (i.e., not breaking up the analysis into scaffold sets), the original *τ*-RAMD simulations showed ranking performance (*ρ* = 0.67 for 70 ligands) that was also poorer than the MW ranking baseline (*ρ*_MW_ = 0.76 for 70 ligands),^10^ highlighting the inherent difficulty in modeling this dataset, even for computationally expensive MD-based protocols. Our ranking performance for the 6 datasets presented in Figure 2(a) outperforms the ligand molecular weight baseline for the other 5 targets studied however (see Figure 2(a) and Table S1). Average simulation run times per complex for the validation targets presented in Figure 2(a) are shown in Figure 2(b). The average run times across all datasets in this validation exercise (71 seconds) represent a reduction in computational cost of at least 3 − 5 orders of magnitude compared to state-of-the-art MD-based alternatives when considering the number of required force field evaluations.

### Predicting Binding Site Mutation Effects on Ligand Unbinding Kinetics

We were also interested in establishing whether our fast unbinding kinetics algorithm could be applied to the related problem of predicting binding site mutation effects upon ligand unbinding kinetics. Such predictions have important applications for areas such as ligand design, protein engineering and personalized medicine. We performed unbinding kinetics simulations on tiotropium interacting with mutated forms of the M_3_ muscarinic receptor (M_3_) to test our simulation methodology’s sensitivity to small changes in the binding site. Tautermann and co-workers previously rationalized experimental ligand RTs using microsecond-scale MD simulations of the bound state. ^32^ More recently, Coricello and co-workers have used adiabatic-bias MD simulations to investigate active site mutants reported for M_3_.^14^ By using path collective variables (PCVs) and running simulations to model the ligand unbinding process from the orthosteric site to the vestibule site (i.e., not full ligand dissociation), they improved computational efficiency, however running a set of 12 M_3_ mutants required 182 GPU days on a NVIDIA GeForce RTX 3060 Ti GPU.^14^ Our results for the M_3_ mutant dataset are shown in Figure 2(c), where we show results for 13 M_3_ mutants, including the 12 mutants reported by Coricello and co-workers. ^14^ Our results show strong predictive accuracy with a computational cost some 5 orders of magnitude lower per mutant complex (Coricello and co-workers reported their predictions with *ρ* = 0.72 and *R*^2^ = 0.48). The M_3_ mutant experimental off rates cover approximately five orders of magnitude yet our energy space method is able to effectively model the emergence of this large dynamical domain from small chemical modifications within the M_3_ binding site.

### Identifying Long-Acting Ligands with the Unbinding Kinetics Simulation Method

To assess the applicability of our method to a challenging kinetics screening task, we combined several unbinding kinetics datasets into a single, larger-scale exercise. This involves the six datasets reported in Figure 2(a), as well as confidential propitiatory datasets belonging to Delphia Therapeutics from an active drug discovery project (discussed in detail later in this work). Our aim is to test what level of enrichment is obtained when classifying *non-transient binders* and *long-acting binders* (defined here as those compounds with residence times greater than one minute and one hour, respectively) across different protein targets. The overall dataset consists of 248 ligand-protein complexes spanning structures from 8 different protein targets. Protein families represented in this exercise include kinases (FAK and TTK), a metaloenzyme (Thermolysin), a GPCR (M_3_), a molecular chaperone protein (HSP90), a cap-binding protein (eIF4E), and the redacted target from the collaboration between scientists at Kvantify and Delphia Therapeutics. The ligands cover a wide range of chemical space, spanning from larger fragments to macromolecules (molecular weights ranging from 283 to 2108 Da with an average molecular weight of 505 Da). The structures of the complexes studied here include crystallographic, docking and ligand alignment examples. Experimental residence time data was principally collected using surface plasmon resonance experiments. For clarity, in this section we convert p*k*_off_ values to their corresponding RTs as

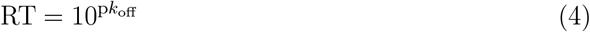

Figure 3(a) shows the unbinding kinetics scores as a function of experimental ligand RTs for the 248 complexes studied. It is important to note that the unbinding kinetics scores computed using our method are not rescaled in any way across protein targets; they are simply the raw output of each independent unbinding kinetics simulation. Despite simultaneously deploying our fast unbinding kinetics simulation method across different highly diverse protein families and ligand types, as well as different experimental structure and kinetics data sources, we obtained a strong overall linear correlation (*R*^2^ = 0.60) and ranking performance (*ρ* = 0.81) that underlines the predictive utility of our methodology. Furthermore, this exercise demonstrates that our methodology is not biased by ligand molecular weight. When considering ligand RTs against ligand molecular weight, we find these quantities to be essentially uncorrelated across the dataset (*R*^2^ = 0.01, *ρ* = −0.11). Even when removing the high molecular weight eIF4E dataset, the same conclusion is reached (*R*^2^ = 0.01, *ρ* = −0.10).

**Figure 3:**
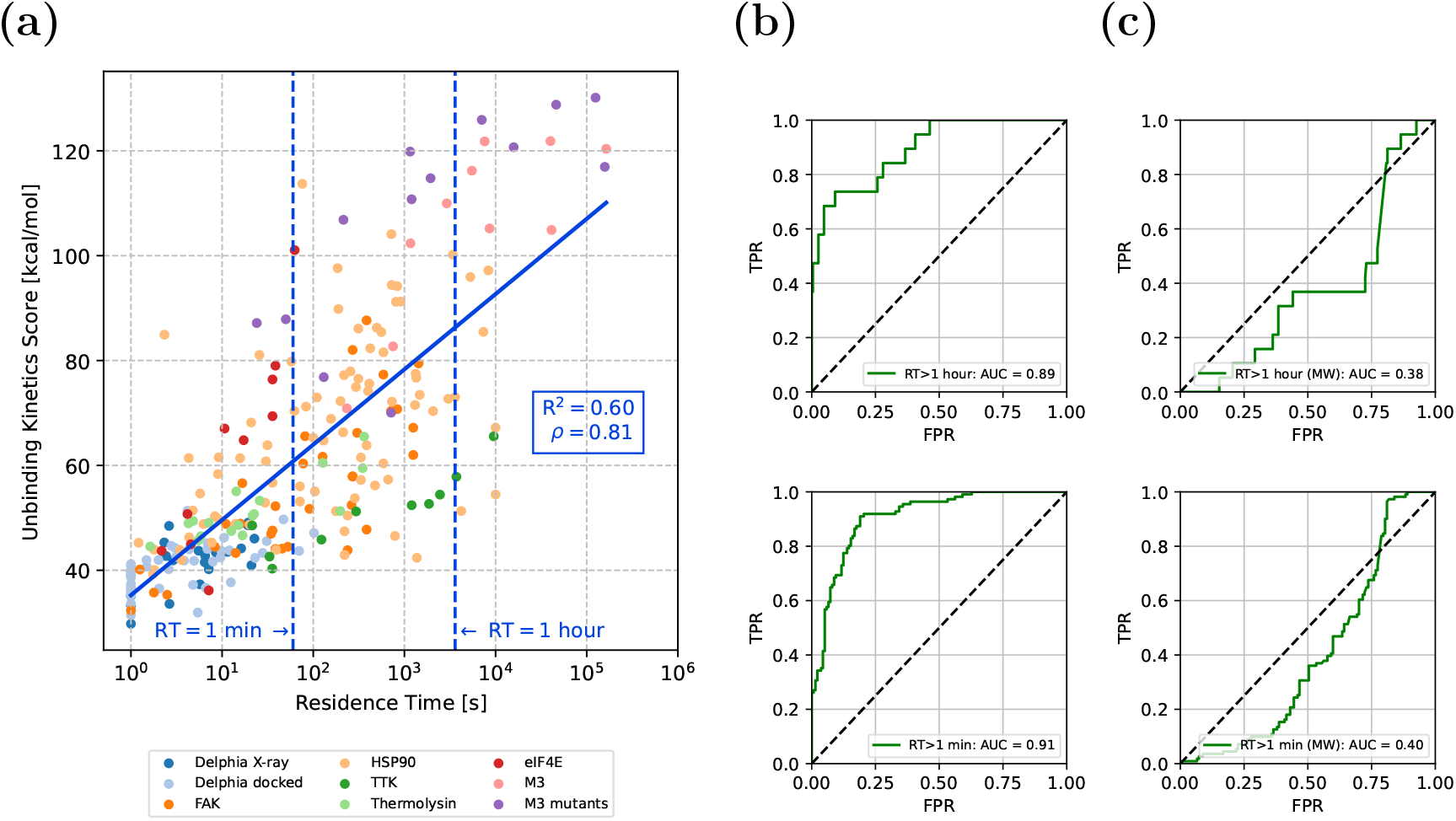
*Koffee Unbinding Kinetics* predicts ligand-protein residence times across different protein targets without rescaling. (a) Scatter plot of unbinding kinetics score as a function of experimental RT measurements. RTs of one second and one hour are indicated in the plot by the dashed vertical lines. (b) Receiver operating characteristic (ROC) curves for both long-acting (RT > one hour, upper) and non-transient (RT > one minute, lower) binder classification using our unbinding kinetics score. (c) ROC curves for both long-acting (RT > one hour, upper) and non-transient (RT > one minute, lower) binder classification using the baseline ligand molecular weight descriptor.

Receiver operating characteristic (ROC) curves for both the long-acting and non-transient binder classification tasks (see Figure 3(b)) yielded AUC-ROC values of 0.89 and 0.91, respectively, indicating strong performance in each of these designated binary classification tasks. The same tasks - using ligand molecular weight as a baseline descriptor - exhibit worse than random performance in both instances (i.e., AUC-ROC < 0.5, see Figure 3(c)). The powerful predictive utility of our method across different protein targets is highlighted by its performance on this broad and challenging exercise. Furthermore, the results from the unbinding kinetics method have been shown to be uncorrelated to the underlying ligand chemistry and molecular weight distribution, a valuable feature of our approach when considering application to real-world drug discovery challenges where modeling highly non-congeneric ligand sets is a common challenge.

### Application of the Unbinding Kinetics Simulation Method to an Active Drug Discovery Program

In collaboration with scientists at Delphia Therapeutics, we used unbinding kinetics simulations to provide ligand unbinding kinetics scoring as part of an active drug discovery program. While the structure-activity relationship (SAR) scheme for the program was well-characterized, the corresponding structure-kinetics relationship (SKR) was initially more ambiguous. The proprietary dataset from the program consists of 20 high-quality X-ray crystal structures, each in complex with a different modulator. These 20 experimentally-derived structures then served as templates for docking for a further 84 compounds that make up the *Docking* (*N* = 49) and *Prospective* (*N* = 35) ligand sets. For each compound, corresponding kinetic measurements were obtained by SPR. In cases where ligand RTs were too short-lived to measure experimentally, a p*k*_off_ value of 0 was assigned. Docking was performed using a core-constrained approach, leveraging the high-quality X-ray structures as templates for additional ligands. We then ran the unbinding kinetics simulation protocol on the *X-ray, Docking* and *Prospective* sets. Figure 4 shows unlabeled results for this project, where we observed excellent discrimination between transient/non-binders (shown in orange) and long-acting binders (shown in green), as well as strong predictive ranking performance on the non-transient binders across the three sets, with *ρ* ∈ (0.54, 0.67). We were interested in two main features of our predictions. Firstly, we were interested to see whether we could accurately identify non-transient (p*k*_off_ > 0) binders. For the three sets investigated, we obtained AUC-ROC scores of 0.98 (*X-ray*), 0.88 (*Docking*) and 0.83 (*Prospective*), indicative of strong enrichment. We also investigated whether we could enrich for ligands in the *Prospective* set with RTs greater than those previously measured in the *X-ray* and *Docking* sets. In this exercise, we chose to define long-acting binders as ligands with p*k*_off_ > 2 (i.e., a RT > 100 s). It was unclear whether such long-acting binders would be present in the *Prospective* set however we obtained an AUC-ROC score of 0.97 for this task, highlighting the ability of our methodology to identify long-acting chemical species unseen in the previous cycles of the project.

**Figure 4:**
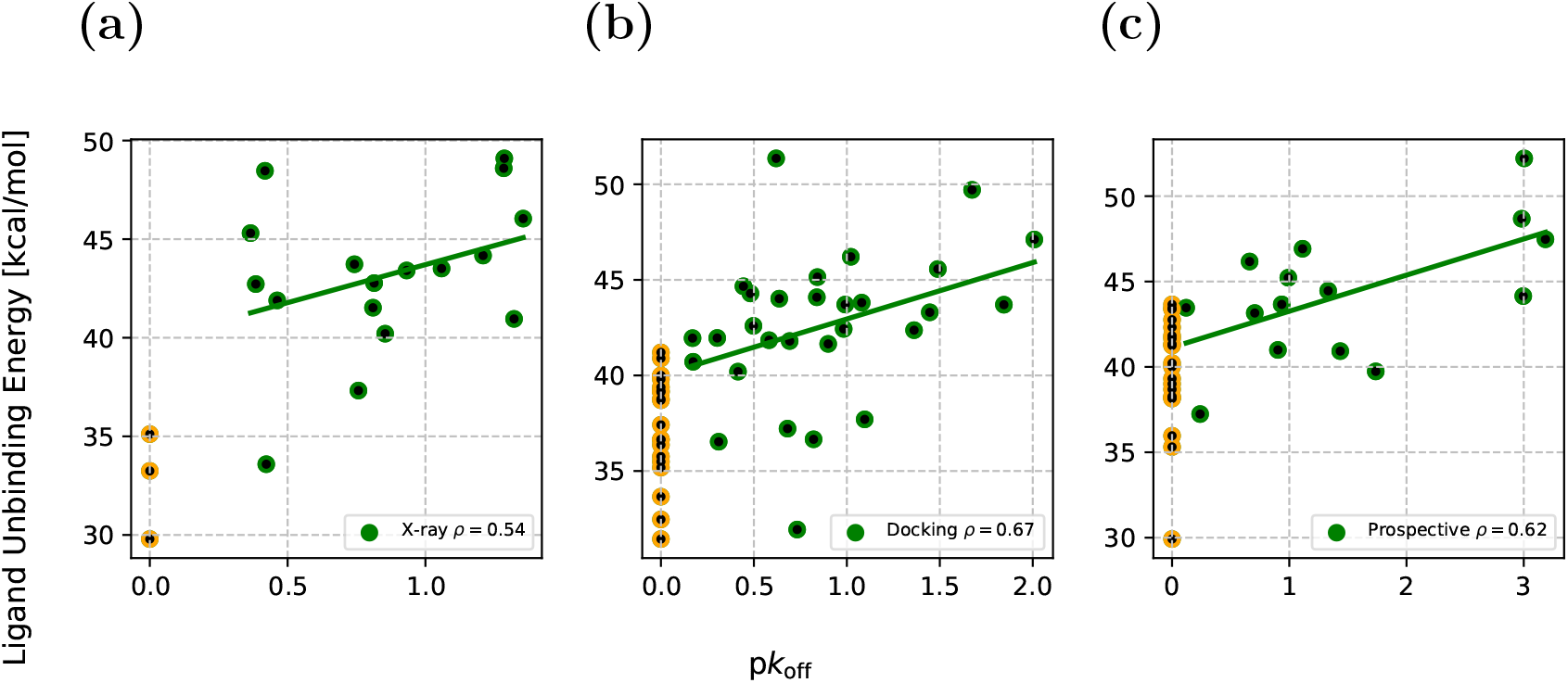
*Koffee Unbinding Kinetics* prospectively identified long-acting binders in an active drug discovery project. Unbinding kinetics predictions for datasets provided by Delphia Therapeutics with transient binders (orange markers) and non-transient binding ligands (green markers). (a) The *X-ray* set contains X-ray crystal structures with previously characterized p*k*_off_ values. (b) The *Docking* set contains ligands poses generated by docking with previously characterized p*k*_off_ values. (c) The *Prospective* set contains further ligand poses generated by docking without previously characterized p*k*_off_ values, which were collected following unbinding kinetics predictions being made. Spearman *ρ* values are calculated for the non-transient binding ligands.

## Discussion

Computer simulations of ligand RTs have previously been limited to low-throughput approaches based on computationally expensive MD simulations, which are also associated with high setup latency costs and considerable user training time investments. As such, these approaches do not suit typical drug discovery project requirements and their real-world deployment has been understandably limited. As a solution to the RT problem, we have developed the *Koffee Unbinding Kinetics* simulation algorithm. In this work, we have shown our simulation methodology provides fast, reliable RT predictions that are applicable across a diverse range of different ligand-protein complexes, including those containing challenging non-congeneric ligand series. Furthermore, we have shown that our methodology is insensitive to ligand molecular weight as well as deploying it on a live drug discovery project. This demonstrated the utility of our unbinding kinetics approach in supporting decision making in a real-life drug discovery environment, where unbinding kinetics scores can be used prospectively to accurately prioritize compounds with desirable RT characteristics for subsequent synthesis and testing.

While advances in hardware mean that MD simulations of ever-increasing timescales and complexity can be performed, on-demand chemical libraries are growing much faster than available computing power. This necessitates the development of entirely new approaches to accurately score these vast number of available compounds to make better use of this ever-expanding chemical space. Our methodology significantly accelerates the calculation of ligand RTs by at least 3 − 5 orders of magnitude compared to MD-based techniques while offering powerful predictive performance. Each simulation is completed in approximately one minute on single NVIDIA T4 GPU with no manual setup required, making *Koffee Unbinding Kinetics* well-suited for rapid deployment to high-throughput workflows in computational drug discovery.

## Acknowledgement

The authors thank Christina Song Levisen for proofreading the manuscript. No reuse of any part of this article is permitted without prior permission from the authors.

## Supporting Information Available

Chemical structures of ligands used in the initial validation exercise; molecular weight dependence of kinetics measurements for the datasets used in the initial validation exercise (.pdf). Details of the construction of FAK and TTK datasets used in this work (.csv).

## References

(1) Copeland, R. A. The drug–target residence time model: a 10-year retrospective. Nature Reviews Drug Discovery 2016, 15, 87–95.

(2) Bradshaw, J. M. et al. Prolonged and tunable residence time using reversible covalent kinase inhibitors. Nature Chemical Biology 2015, 11, 525–531.

(3) Bernetti, M.; Masetti, M.; Rocchia, W.; Cavalli, A. Kinetics of Drug Binding and Residence Time. Annual Review of Physical Chemistry 2019, 70, 143–171.

(4) Waring, M. J.; Arrowsmith, J.; Leach, A. R.; Leeson, P. D.; Mandrell, S.; Owen, R. M.; Pairaudeau, G.; Pennie, W. D.; Pickett, S. D.; Wang, J.; Wallace, O.; Weir, A. An analysis of the attrition of drug candidates from four major pharmaceutical companies. Nature Reviews Drug Discovery 2015, 14, 475–486.

(5) Guo, D.; Mulder-Krieger, T.; IJzerman, A. P.; Heitman, L. H. Functional efficacy of adenosine A2A receptor agonists is positively correlated to their receptor residence time. British Journal of Pharmacology 2012, 166, 1846–1859.

(6) Kordylewski, S. K.; Bugno, R.; Podlewska, S. Residence time in drug discovery: current insights and future perspectives. Pharmacological Reports 2025, 77, 851–873.

(7) Bose, S.; Lotz, S. D.; Deb, I.; Shuck, M.; Lee, K. S. S.; Dickson, A. How Robust Is the Ligand Binding Transition State? Journal of the American Chemical Society 2023, 145, 25318–25331.

(8) Lee, E. H.; Hsin, J.; Sotomayor, M.; Comellas, G.; Schulten, K. Discovery Through the Computational Microscope. Structure 2009, 17, 1295–1306.

(9) Zia, S. R.; Coricello, A.; Bottegoni, G. Increased throughput in methods for simulating protein ligand binding and unbinding. Current Opinion in Structural Biology 2024, 87, 102871.

(10) Kokh, D. B.; Amaral, M.; Bomke, J.; Grädler, U.; Musil, D.; Buchstaller, H.-P.; Dreyer, M. K.; Frech, M.; Lowinski, M.; Vallee, F.; Bianciotto, M.; Rak, A.; Wade, R. C. Estimation of Drug-Target Residence Times by -Random Acceleration Molecular Dynamics Simulations. Journal of Chemical Theory and Computation 2018, 14, 3859– 3869.

(11) Sinko, W.; Mertz, B.; Shimizu, T.; Takahashi, T.; Terada, Y.; Kimura, S. R. ModBind, a Rapid Simulation-Based Predictor of Ligand Binding and Off-Rates. Journal of Chemical Information and Modeling 2025, 65, 265–274.

(12) Ojha, A. A.; Votapka, L. W.; Amaro, R. E. QMrebind: incorporating quantum mechanical force field reparameterization at the ligand binding site for improved drug-target kinetics through milestoning simulations. Chemical Science 2023, 14, 13159–13175.

(13) Ojha, A. A.; Votapka, L. W.; Amaro, R. E. Advances and Challenges in Milestoning Simulations for Drug–Target Kinetics. Journal of Chemical Theory and Computation 2024, 20, 9759–9769.

(14) Coricello, A.; Chiaravalle, A. L.; Musgaard, M.; Tehan, B. G.; Elisi, G. M.; Bottegoni, G. Adiabatic-Bias Molecular Dynamics Simulations Reveal the Impact of Mutations on Muscarinic Antagonist Unbinding Kinetics. Journal of Chemical Information and Modeling 2025, 65, 7129–7142.

(15) Lee, S.; Wang, D.; Seeliger, M. A.; Tiwary, P. Calculating Protein–Ligand Residence Times through State Predictive Information Bottleneck Based Enhanced Sampling. Journal of Chemical Theory and Computation 2024, 20, 6341–6349.

(16) Smith, Z.; Branduardi, D.; Lupyan, D.; D’Arrigo, G.; Tiwary, P.; Wang, L.; Krilov, G. Towards automated physics-based absolute drug residence time predictions. ChemRxiv 2025, doi:10.26434/chemrxiv-2025-wg75c.

(17) Serrano-Morrás, A.; Westermaier, Y.; Majewski, M.; Barril, X. The Quasi-Bound State as a Predictor of Relative Binding Free Energy. Journal of Chemical Information and Modeling 2025, 65, 5544–5552.

(18) Maier, J. A.; Martinez, C.; Kasavajhala, K.; Wickstrom, L.; Hauser, K. E.; Simmerling, C. ff14SB: Improving the Accuracy of Protein Side Chain and Backbone Parameters from ff99SB. Journal of Chemical Theory and Computation 2015, 11, 3696–3713.

(19) Wang, J.; Wolf, R. M.; Caldwell, J. W.; Kollman, P. A.; Case, D. A. Development and testing of a general amber force field. Journal of Computational Chemistry 2004, 25, 1157–1174.

(20) Wang, J.; Wang, W.; Kollman, P. A.; Case, D. A. Automatic atom type and bond type perception in molecular mechanical calculations. Journal of Molecular Graphics and Modelling 2006, 25, 247–260.

(21) Mongan, J.; Simmerling, C.; McCammon, J. A.; Case, D. A.; Onufriev, A. Generalized Born Model with a Simple, Robust Molecular Volume Correction. Journal of Chemical Theory and Computation 2007, 3, 156–169.

(22) Nguyen, H.; Roe, D. R.; Simmerling, C. Improved Generalized Born Solvent Model Parameters for Protein Simulations. Journal of Chemical Theory and Computation 2013, 9, 2020–2034.

(23) Tomić, A.; Horvat, G.; Ramek, M.; Agić, D.; Brkić, H.; Tomić, S. New Zinc Ion Parameters Suitable for Classical MD Simulations of Zinc Metallopeptidases. Journal of Chemical Information and Modeling 2019, 59, 3437–3453.

(24) Raguette, L. E.; Cuomo, A. E.; Belfon, K. A. A.; Tian, C.; Hazoglou, V.; Witek, G.; Telehany, S. M.; Wu, Q.; Simmerling, C. phosaa14SB and phosaa19SB: Updated Amber Force Field Parameters for Phosphorylated Amino Acids. Journal of Chemical Theory and Computation 2024, 20, 7199–7209.

(25) Eastman, P. et al. OpenMM 8: Molecular Dynamics Simulation with Machine Learning Potentials. The Journal of Physical Chemistry B 2024, 128, 109–116.

(26) Schuetz, D. A. et al. Kinetics for Drug Discovery: an industry-driven effort to target drug residence time. Drug Discovery Today 2017, 22, 896–911.

(27) Liu, H.; Su, M.; Lin, H.-X.; Wang, R.; Li, Y. Public Data Set of Protein-Ligand Dissociation Kinetic Constants for Quantitative Structure–Kinetics Relationship Studies. ACS Omega 2022, 7, 18985–18996.

(28) Lama, D.; Liberatore, A.-M.; Frosi, Y.; Nakhle, J.; Tsomaia, N.; Bashir, T.; Lane, D. P.; Brown, C. J.; Verma, C. S.; Auvin, S. Structural insights reveal a recognition feature for tailoring hydrocarbon stapled-peptides against the eukaryotic translation initiation factor 4E protein. Chemical Science 2019, 10, 2489–2500.

(29) Galvani, F.; Pala, D.; Cuzzolin, A.; Scalvini, L.; Lodola, A.; Mor, M.; Rizzi, A. Unbinding Kinetics of Muscarinic M3 Receptor Antagonists Explained by Metadynamics Simulations. Journal of Chemical Information and Modeling 2023, 63, 2842–2856.

(30) Uitdehaag, J. C.; de Man, J.; Willemsen-Seegers, N.; Prinsen, M. B.; Libouban, M. A.; Sterrenburg, J. G.; de Wit, J. J.; de Vetter, J. R.; de Roos, J. A.; Buijsman, R. C.; Zaman, G. J. Target Residence Time-Guided Optimization on TTK Kinase Results in Inhibitors with Potent Anti-Proliferative Activity. Journal of Molecular Biology 2017, 429, 2211–2230.

(31) Wang, R.; Fang, X.; Lu, Y.; Wang, S. The PDBbind Database: Collection of Binding Affinities for Protein-Ligand Complexes with Known Three-Dimensional Structures. Journal of Medicinal Chemistry 2004, 47, 2977–2980, PMID: 15163179.

(32) Tautermann, C. S.; Kiechle, T.; Seeliger, D.; Diehl, S.; Wex, E.; Banholzer, R.; Gantner, F.; Pieper, M. P.; Casarosa, P. Molecular Basis for the Long Duration of Action and Kinetic Selectivity of Tiotropium for the Muscarinic M3 Receptor. Journal of Medicinal Chemistry 2013, 56, 8746–8756.

(33) Eastman, P.; Swails, J.; Chodera, J. D.; McGibbon, R. T.; Zhao, Y.; Beauchamp, K. A.; Wang, L.-P.; Simmonett, A. C.; Harrigan, M. P.; Stern, C. D.; Wiewiora, R. P.; Brooks, B. R.; Pande, V. S. OpenMM 7: Rapid development of high performance algorithms for molecular dynamics. PLOS Computational Biology 2017, 13, 1–17.

(34) Bietz, S.; Urbaczek, S.; Schulz, B.; Rarey, M. Protoss: a holistic approach to predict tautomers and protonation states in protein-ligand complexes. Journal of Cheminformatics 2014, 6, 12.

(35) López-Pérez, K.; Kim, T. D.; Miranda-Quintana, R. A. iSIM: instant similarity. Digital Discovery 2024, 3, 1160–1171.

(36) Xu, C.; Zhang, X.; Zhao, L.; Verkhivker, G. M.; Bai, F. Accurate Characterization of Binding Kinetics and Allosteric Mechanisms for the HSP90 Chaperone Inhibitors Using AI-Augmented Integrative Biophysical Studies. JACS Au 2024, 4, 1632–1645.

