## Supplementary material for "Unlocking Scalable Ligand Residence Time Predictions with Koffee Unbinding Kinetics Simulations": SI.pdf

### Supporting Information for “Unlocking Scalable Ligand Residence Time Predictions with Koffee Unbinding Kinetics Simulations”

Niels Kristian Madsen,[*,†,‡] Robert M. Ziolek,[*,†,‡] Daniel Kongsgaard,[†] Christian Flohr Nielsen,[†] Anders Dyhr Nørløv,[†] Daniela Dolciami,[†] Joshua R. Sacher,[¶] Klaus Michelsen,[¶] Michael G. Acker,[¶] Nils Anton Berglund,[†] Mikael H. Christensen,[†] Allan Grønlund,[†] Lise Husted,[†] David E. Gloriam,[†,§] Albert J. Kooistra,[†,§] and Nikolaj Thomas Zinner[†]

†*Kvantify ApS, 2100 Copenhagen, Denmark*

‡*N.K.M. and R.M.Z. contributed equally*

¶*Delphia Therapeutics, 1400 One Kendall Square, Cambridge, Massachusetts 02139, United States*

§*Department of Drug Design and Pharmacology, Faculty of Health and Medical Sciences, University of Copenhagen, 2100 Copenhagen, Denmark*

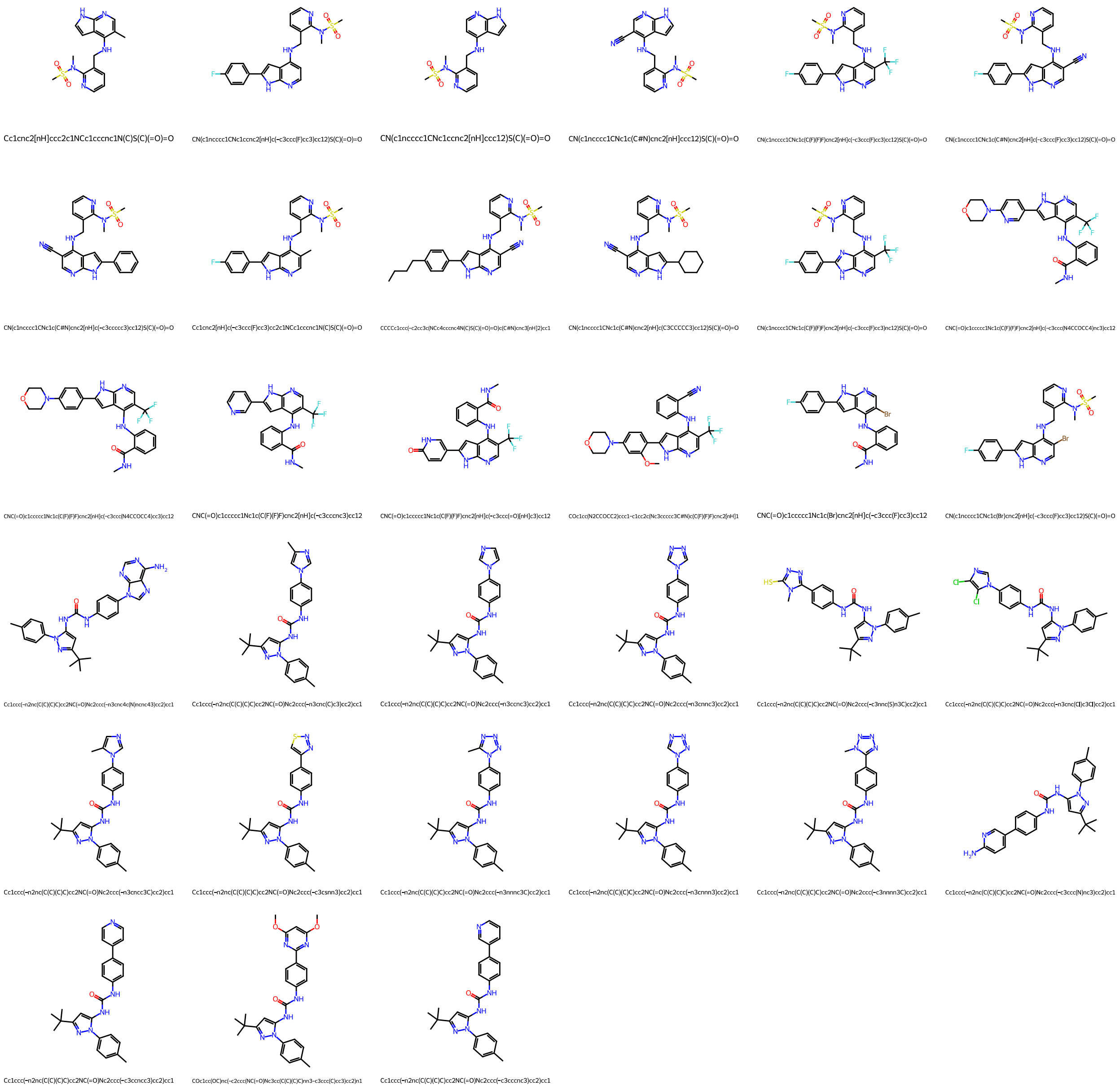


Figure S1: Chemical structures of the FAK ligand set.

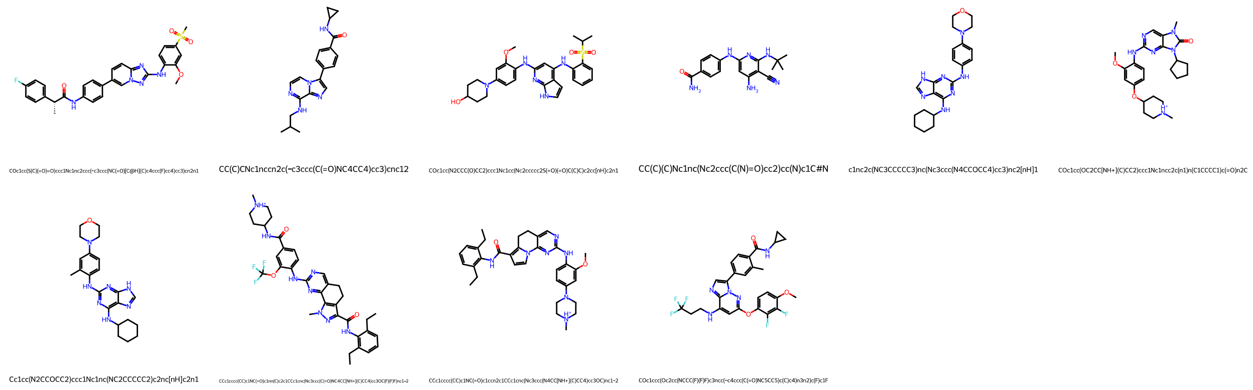


Figure S2: Chemical structures of the TTK ligand set.

Table S1: Spearman correlation between ligand molecular weight and experimentally measured $pk_{\text{off}}$ ($\rho_{\text{MW}}$).

| Target | $\rho_{\text{MW}}$ |
|---|---|
| eIF4E | 0.83 |
| FAK | 0.22 |
| HSP90 | 0.76 |
| M3 | 0.32 |
| M3 mutants | N/A |
| Thermolysin | 0.74 |
| TTK | 0.52 |
